# Membrane metalloendopeptidase suppresses prostate carcinogenesis by attenuating effects of gastrin-releasing peptide on stem/progenitor cells

**DOI:** 10.1101/2019.12.23.887356

**Authors:** Chieh-Yang Cheng, Zongxiang Zhou, Meredith Stone, Bao Lu, Andrea Flesken-Nikitin, David M. Nanus, Alexander Yu. Nikitin

## Abstract

Aberrant neuroendocrine signaling is frequent yet poorly understood feature of prostate cancers. Membrane metalloendopeptidase (MME) is responsible for the catalytic inactivation of neuropeptide substrates, and is downregulated in nearly 50% of prostate cancers. However its role in prostate carcinogenesis, including formation of castration-resistant prostate carcinomas, remains uncertain. Here we report that MME cooperates with PTEN in suppression of carcinogenesis by controlling activities of prostate stem/progenitor cells. Lack of MME and PTEN results in development of adenocarcinomas characterized by propensity for vascular invasion and formation of proliferative neuroendocrine clusters after castration. Effects of MME on prostate stem/progenitor cells depend on its catalytic activity and can be recapitulated by addition of the MME substrate, gastrin-releasing peptide (GRP). Knockdown or inhibition of GRP receptor (GRPR) abrogate effects of MME deficiency, and delay growth of human prostate cancer xenografts by reducing the number of cancer propagating cells. In sum, our study provides a definitive proof of tumor suppressive role of MME, links GRP/GRPR signaling to the control of prostate stem/progenitor cells, and shows how dysregulation of such signaling may promote formation of castration-resistant prostate carcinomas. It also identifies GRPR as a valuable target for therapies aimed at eradication of cancer propagating cells in prostate cancers with MME downregulation.

## Introduction

Prostate cancer is the most frequently diagnosed cancer and is the second leading cause of cancer-related death in men in the United States ^38^. While most prostate cancers are adenocarcinomas, a significant percentage also have dysregulation of neuroendocrine signaling, such as excessive accumulation of cells with neuroendocrine differentiation and/or overproduction of neuropeptides ^8, 19, 27^. A large amount of data demonstrate neuropeptides, such as gastrin-releasing peptide (GRP), are associated with accelerated prostate cancer progression and inferior prognosis ^34, 45, 48^. GRP can promote cell proliferation, and accelerate migration and invasion of prostate cancer cells ^15, 34, 41, 54^. Targeting of the GRP receptor suppresses growth in cell culture and xenograft models ^26^. However, specific mechanisms by which neuropeptide dysregulation contributes to the pathogenesis of prostate cancer remain insufficiently elucidated.

Local concentration of neuropeptides is regulated in part by membrane metallo-endopeptidase (MME, aka Neutral endopeptidase). MME is a cell-surface peptidase member of the M13 family of zinc peptidases, which also includes endothelin converting enzymes (ECE-1 and ECE-2), KELL and PEX. MME cleaves peptide bonds on the amino side of hydrophobic amino acids and is the key enzyme in processing of a variety of physiologically active peptides, such as GRP, neurotensin, and vasoactive intestinal peptide ^10, 54^. MME is downregulated in nearly 50% of primary and metastatic prostate cancers, independently predicting an inferior prognosis ^12, 17, 30^. In addition to its downregulation by androgen withdrawal ^37, 55^, MME expression is also downregulated by methylation, suggesting its tumor suppressive effects ^30, 49^. Indeed, MME expression reduces growth, motility ^41^, and survival ^40, 42^ of prostate cancer cells in cell culture. Consistent with these observations, replacements of MME inhibit tumorigenicity of prostate cancer cells in xenograft experiments ^7, 14^. Nevertheless, mice lacking *Mme* show no prostate cancer-related phenotype ^24^, and the role of MME in prostate cancer progression remains uncertain.

At least part of MME effects are mediated by the PI3K/AKT pathway that plays a key role in multiple cellular processes including cell survival, proliferation, and cell migration reviewed in ^44^. MME associates with and stabilizes the PTEN tumor suppressor protein, resulting in increased PTEN phosphatase activity, thereby inhibiting AKT activating phosphorylation ^43^. MME may also have PTEN independent mechanisms of AKT inhibition by processing neuropeptides, such as GRP, which are known to activate AKT ^42^. Consistent with a possibility of potential cooperation between MME and PTEN in suppression of carcinogenesis, downregulation of MME is observed in 42% and 63% of PTEN deficient cases of human primary and metastatic prostate cancers, respectively^46^. However, it remains unknown if catalytically dependent neuropeptide-based mechanisms of MME tumor suppression play a role in prostate cancer progression.

The mouse prostate is composed of a series of branching ducts, each containing distal and proximal regions relative to the urethra ^39^. Proliferating, transit-amplifying cells are preferentially located in the distal region of the prostatic ducts, whereas cells with stem cell-like properties, such as low cycling rate, self-renewal ability, high *ex vivo* proliferative potential, and androgen withdrawal resistance, mainly reside in the proximal region of the prostatic ducts ^5, 36, 47, 57,Fu, 2018 #17167^. Thus, approaches based on the isolation of cells according to their displayed stem cell-specific markers can be complemented by careful evaluation of stem cell compartments *in situ*.

In the current study, we used autochthonous mouse model of prostate neoplasia associated with deficiency of *Pten* tumor suppressor gene. In this model, prostate carcinogenesis is initiated by the prostate epithelium-specific inactivation of *Pten* driven by PB-*Cre4* transgene (PtenPE-/- mice; ^3, 16, 23, 50, 53, 56^. The majority of mice show early stages of prostate cancer, such as high-grade prostatic intraepithelial neoplasms (HG-PINs) and few animals show early adenocarcinomas characterized by stromal invasion. Thus, it is well suitable for testing if additional genetic alterations, such as *Mme* inactivation, may accelerate cancer progression. We report that lack of both MME and PTEN leads to aggressive prostate cancers manifesting frequent vascular invasion and increased neuroendocrine differentiation after castration. Formation of such cancers is preceded by morphologically detectable neoplastic lesions at the prostate stem/progenitor cell compartment. The effect of MME deficiency on stem/progenitor cells can be recapitulated by its substrate GRP and is abrogated by either GRP receptor (GRPR) antagonist or *GRPR* siRNA knockdown. Knockdown or inhibition of GRP receptor (GRPR) delay growth of human prostate cancer xenografts by reducing the pool of cancer propagating cells.

## Results

### MME cooperates with PTEN in suppression of prostate cancer in autochthonous mouse model

To test the cooperation of *Mme* and *Pten* genes in suppression of prostate cancer in vivo we first evaluated MME expression in HG-PINs and early invasive adenocarcinomas typical for *Pten*^PE-/-^ mice. While irregular MME expression was observed in the majority of neoplastic lesions, MME was absent in the areas of stromal invasion (Fig. 1 and Supplementary Fig. 1). No significant alterations in MME expression were detected in the proximal regions of prostatic ducts, consistent with the lack of neoplastic lesions in that part of the prostate in *Pten*^PE-/-^ mice (Fig. 2a, Supplementary Fig. 2a, and Supplementary Table 1). Consistent with the reported regulation of MME by the androgen receptor (AR) ^29, 37, 55^, castration of both WT and *Pten*^PE-/-^ mice resulted in downregulation but not complete abrogation of MME expression (Fig. 2).

**Fig. 1.**
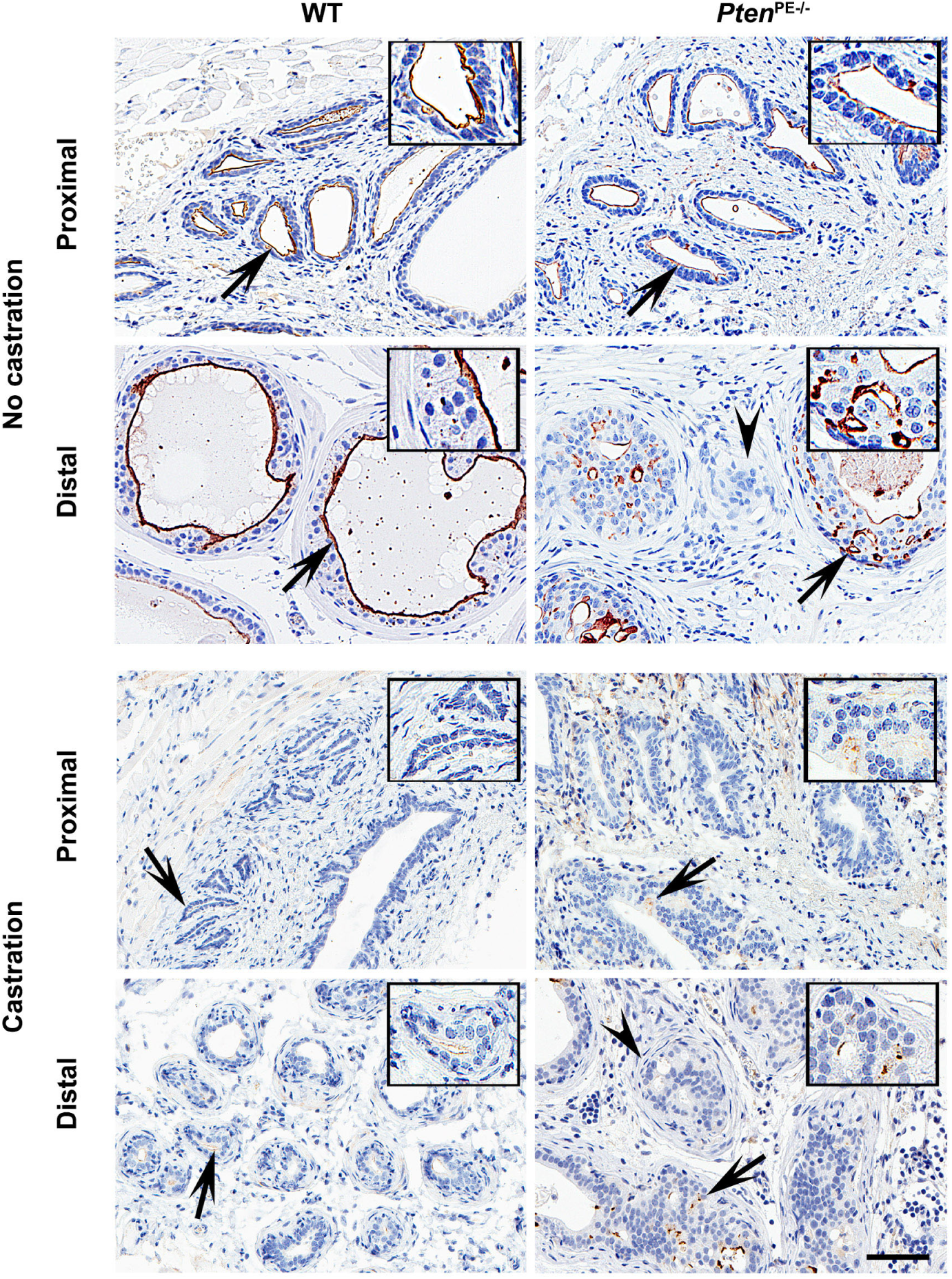
Alterations of *MME* expression in prostate adenocarcinoma of *Pten*^PE-/-^ mice. MME expression in the proximal and distal regions of prostatic ducts of wild-type (WT) and *Pten*^PE-/-^ 12 months old mice, non-castrated and castrated at 5 months of age. Arrows indicate areas shown insets. Note that areas of stromal invasion (arrowheads) by early adenocarcinoma lacks MME expression in *Pten*^PE-/-^ mice. The ABC Elite method with hematoxylin counterstaining was performed. Scale bar, 60 μm and 30 µm (insets). Immunostainings are representative of six WT and six *Pten*^PE-/-^ mice, respectively.

**Fig. 2.**
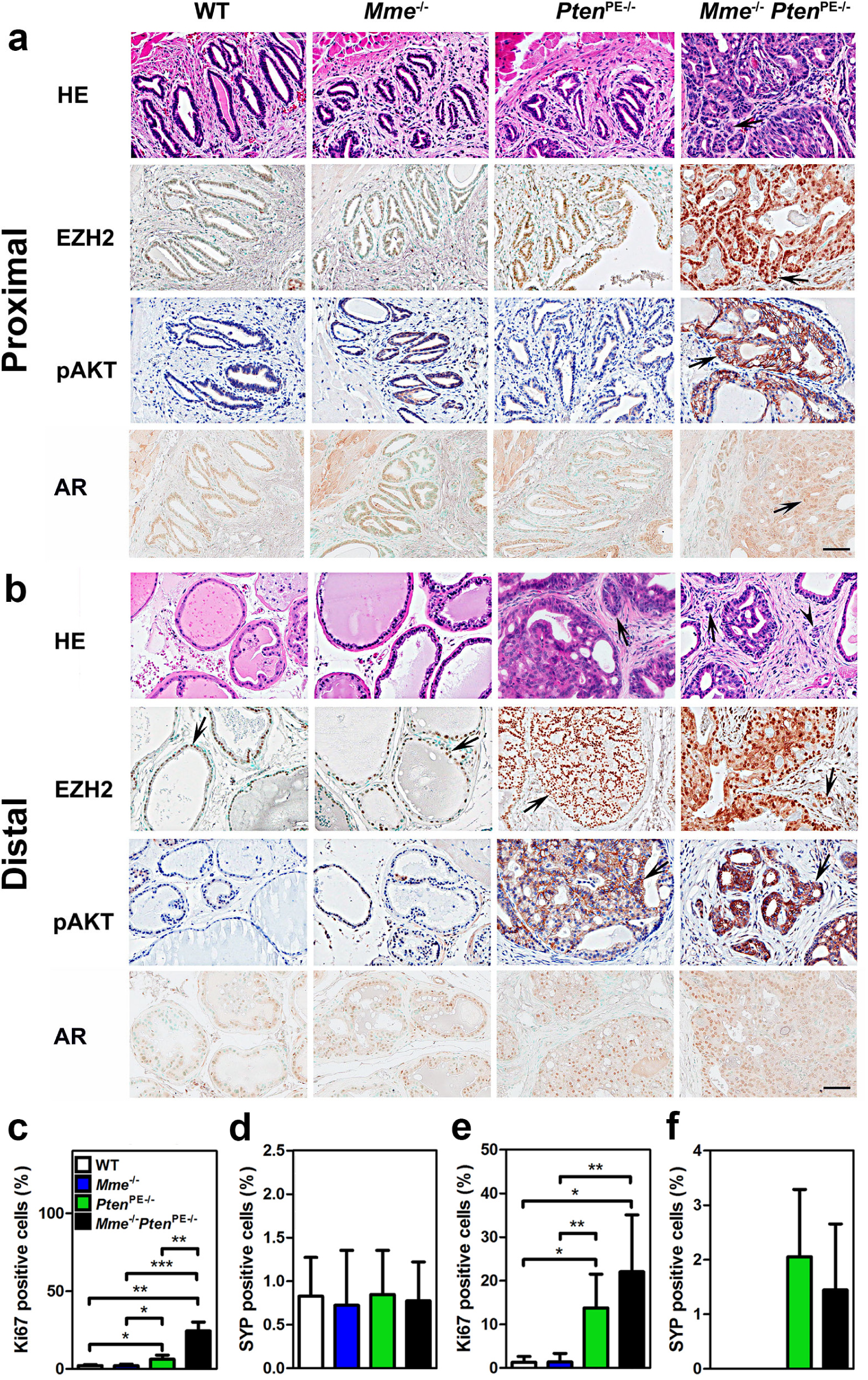
*Mme* and *Pten* cooperate in suppression of prostate carcinogenesis in the proximal regions of prostatic ducts of the mouse. **a-b** Proximal (**a**) and distal (**b**) regions of prostatic ducts in 16-month-old WT (n=6), *Mme^-/-^* (n=15), *Pten*^PE-/-^ (n=14) and *Mme^-/-^Pten*^PE-/-^ (n=15) mice. The arrows indicate stromal invasion of adenocarcinoma (HE) in *Mme^-/-^* and *Mme^-/-^Pten*^PE-/-^ mice and positive immunostained cells. The arrowhead in distal region (HE) indicates vascular invasion in *Mme^-/-^Pten*^PE-/-^ mice. As compared to the prostate epithelium of WT, *Mme^-/-^*, and *Pten*^PE-/-^ mice, adenocarcinomas of *Mme^-/-^Pten*^PE-/-^ mice show higher expression levels of EZH2 and pAKT, but no differences in AR expression. HE, hematoxylin and eosin staining. The ABC Elite method with hematoxylin (pAKT) or methyl green (EZH2, and AR) counterstaining was performed. Scale bar, 60 µm for all images. **b** and **c**. A quantitative analysis of the proliferation rate (**c**, **e**) and frequency of SYP positive cells (**d**, **f**) in proximal (**c**, **d**) and distal (**e**, **f**) regions of prostatic ducts. Annotations for groups in **c-f** are shown in **c**. *P<0.05; **P<0.01; ***P<0.001. Error bars denote SD. All results are representative of six mice per genotype.

Next, we tested if MME deficiency can accelerate prostate carcinogenesis in *Pten*^PE-/-^ mouse model. We crossed *Mme*^-/-^ and *Pten*^PE-/-^ mice, and evaluated prostates of age-matched wild-type (WT), *Mme^-/-^*, *Pten*^PE-/-^ and *Mme^-/-^Pten*^PE-/-^ strains (Fig. 2, Supplementary Fig. 2, and Supplementary Table 1). Consistent with previous observations ^24^, *Mme^-/-^* mice did not develop any neoplastic lesions by 16 months of age, while *Pten*^PE-/-^ mice showed low- and high-grade PINs at 3 months. At 16 months 29% of *Pten*^PE-/-^ mice developed early invasive adenocarcinomas, characterized by separate nests of neoplastic cells in desmoplastic stroma (Fig. 2 and Supplementary Table 1). All neoplastic lesions were in the distal regions of prostatic ducts. Eighty six percent of *Mme^-/-^Pten*^PE-/-^ mice developed adenocarcinomas in the same location (Fisher’s exact test P=0.0025). Furthermore, contrary to *Pten*^PE-/-^ mice, *Mme^-/-^Pten*^PE-/-^ mice developed dysplastic lesions followed by adenocarcinomas in the proximal regions of the prostatic ducts (Fig. 2a, Supplementary Fig. 1 and Supplementary Table 1). Some adenocarcinomas of *Mme^-/-^Pten*^PE-/-^ mice showed distinct vascular invasion, a feature not characteristic for prostatic lesions in *Pten*^PE-/-^ mice (Fig. 2a and Supplementary Fig. 2 and 3). Consistent with these histological observations, prostatic epithelium of *Mme^-/-^Pten*^PE-/-^ mice was characterized by significantly higher proliferative rate according to Ki67 staining (Fig. 2 and Supplementary Fig. 2), expressed higher amounts of epigenetic reprogramming factor EZH2 (Fig. 2 and Supplementary Fig. 4), and showed increased number of CK5 and p63 positive cells (Supplementary Fig. 2) as compared to prostatic lesions in *Pten*^PE-/-^ mice. Prostatic epithelium lesions of *Mme^-/-^Pten*^PE-/-^ mice also showed higher levels of pAKT (Fig. 2 and Supplementary Fig. 4), confirming additional MME-dependent mechanisms of regulation of this downstream target of PTEN ^42, 43^. Consistent with our previous observation ^23^, prostatic neoplastic lesions of *Pten*^PE-/-^ mice had increased number of synaptophysin positive neuroendocrine cells in the distal regions of prostatic ducts (Fig. 2 and Supplementary Fig. 2). However, this number was not additionally elevated in *Mme^-/-^Pten*^PE-/-^ mice. In sum, lack of both MME and PTEN not only promoted lesions typically observed in the prostates of *Pten*^PE-/-^ mice, but also resulted in a distinct new neoplasms located in the proximal regions of prostatic ducts.

### MME loss leads to more aggressive tumor phenotype with increased proliferation of neuroendocrine clusters after castration

As compared to *Pten*^PE-/-^ mice, recurrent tumors in all castrated *Mme^-/-^Pten*^PE-/-^ mice (n=8) had increased levels of phosphorylated MET, and reduced expression of E-cadherin, signatures of more aggressive phenotype (Fig. 3). As previously reported ^23^, recurrent tumors in castrated *Pten*^PE-/-^ mice show decreased expression of AR and increased number of neuroendocrine cells. MME deficiency in castrated *Pten*^PE-/-^ mice did not further affect AR expression levels (Fig. 3). However, recurrent tumors in *Mme^-/-^Pten*^PE-/-^ mice had higher frequency of neuroendocrine clusters (≥ 3 cells) and such clusters were more proliferative according to Ki67 staining (Fig. 3). Thus, MME deficiency promotes the increase in neuroendocrine differentiation of neoplastic cells after castration.

**Fig. 3.**
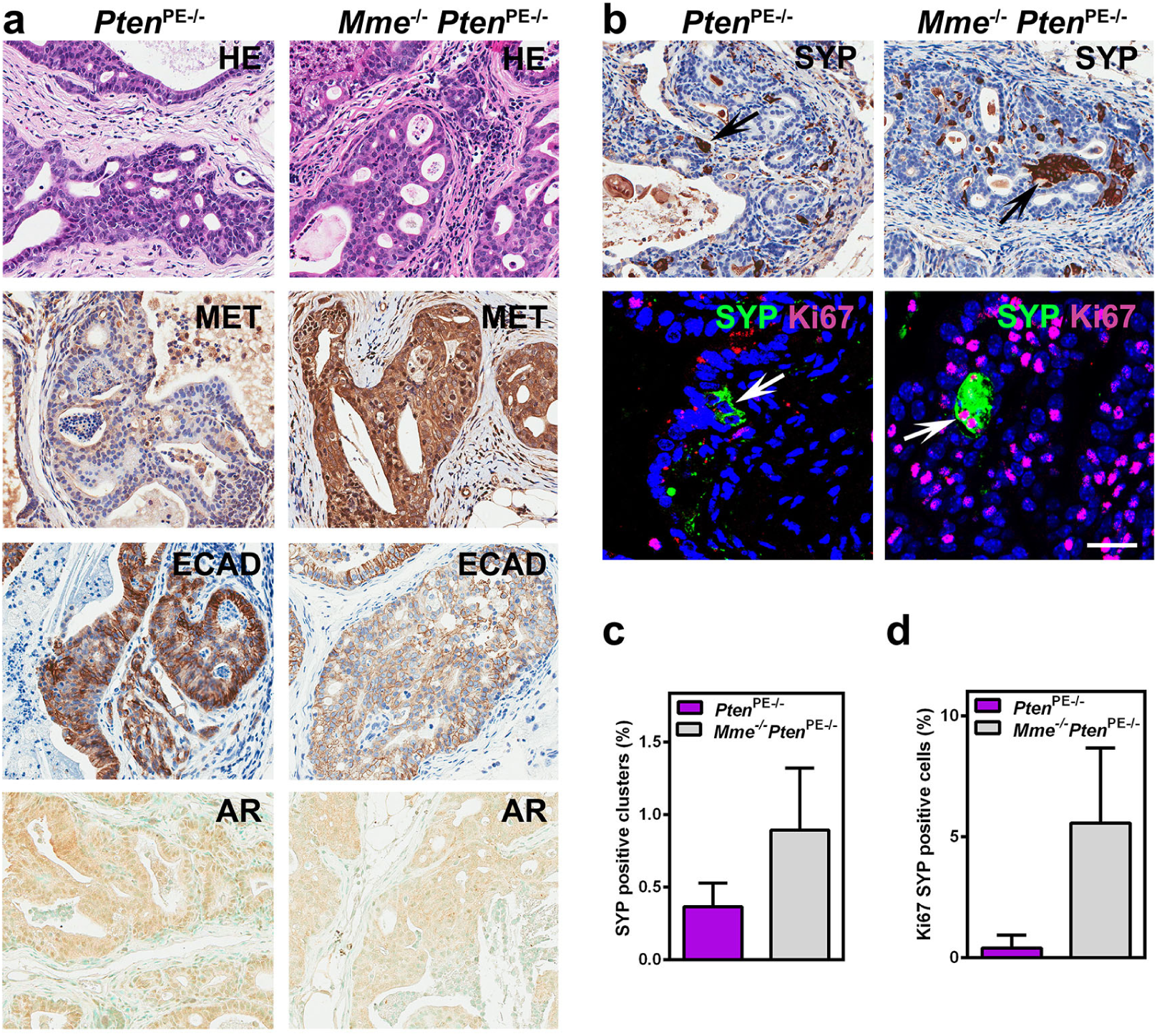
MME deficiency promotes aggressiveness of prostate carcinomas occurring in *Pten*^PE-/-^ cells after castration. **a** Distal regions of prostatic ducts in 16-month-old *Pten*^PE-/-^ (n=6), and *Mme^-/-^Pten*^PE-/-^ (n=8) mice 8 months after castration. The arrows indicate positive immunostained cells. As compared to the prostate epithelium of *Pten*^PE-/-^ mice, adenocarcinomas of *Mme^-/-^Pten*^PE-/-^ mice show higher expression levels of MET, lowered expression of E-Cadherin (ECAD), and no change in AR expression. **b**-**d** Increased number and proliferation rate of SYP positive clusters. Immunostaining (**b**) and quantitative analysis of the frequency of SYP positive clusters (**c**), and SYP cells expressing Ki67 therein (**d**) in distal regions of prostatic ducts. HE, hematoxylin and eosin staining. The ABC Elite method with hematoxylin (MET, ECAD and SYP) or methyl green (AR) counterstaining was performed. Double immunofluorescence with SYP (green) and Ki67 (red) with DAPI (blue) counterstaining. Scale bar, 60 µm for HE and all immunoperoxidase images, and 30 µm for fluorescence images. **c** and **d**, P<0.01. Error bars denote SD. All results are representative of six mice per genotype.

### MME loss promotes activities of PTEN-deficient mouse prostate stem/progenitor cells

The proximal regions of prostatic ducts are particularly enriched in prostate epithelium stem/progenitor cells ^5, 13, 36, 47, 57^. Thus we evaluated potential effects of MME and PTEN deficiency on prostate stem/progenitor cells, isolated as the CD49f^hi^/Sca-1^+^ fraction by fluorescence-activated cell sorting (FACS). *Mme^-/-^* mice had the same number of stem/progenitor cells as age-matched WT mice (5.7% vs 5.6%). In contrast, *Pten*^PE-/-^ mice showed a significant increase of the stem cell pool (8.4%) consistent with previous reports suggesting PTEN’s involvement in regulation of prostate stem/progenitor cells ^9, 28, 51^. The pool of CD49f^hi^/Sca-1^+^ cells deficient for both PTEN and MME, constituted 12.1% of the prostate epithelium, representing an additional 44% increase as compared to PTEN deficient CD49f^hi^/Sca-1^+^ cells (Fig. 4a, P<0.0001).

**Fig. 4.**
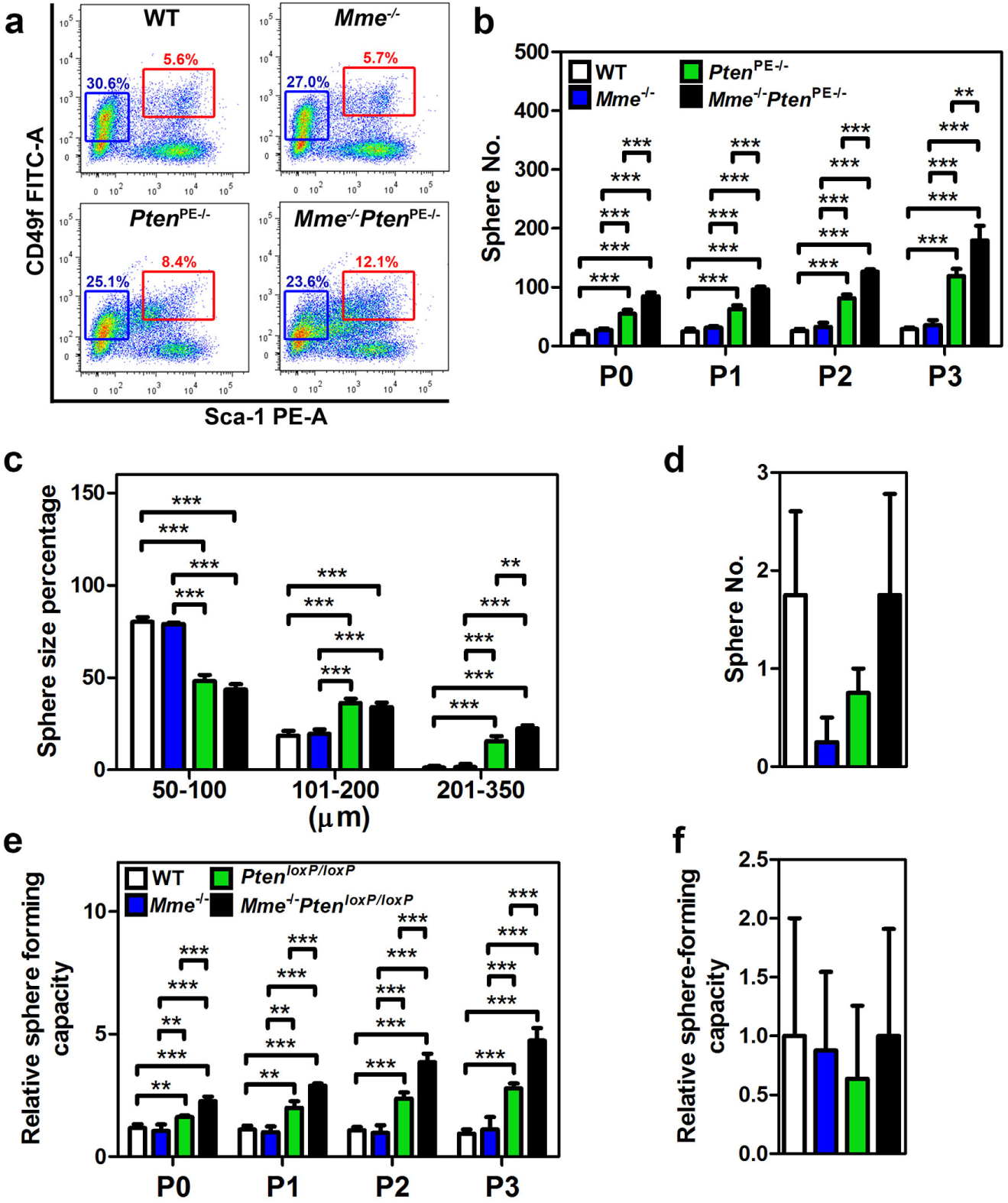
Lack of *Mme* promotes *Pten*-deficient prostate stem/progenitor cell expansion and sphere-forming capacity. **a** Quantitative analysis of distribution of CD49f^hi^/Sca-1^+^ stem/progenitor cells and CD49f^lo^/Sca-1^-^ luminal cells isolated from 3-month-old WT, *Mme^-/-^*, *Pten*^PE-/-^, and *Mme^-/-^Pten*^PE-/-^ mice (n=6 per genotype). Red and blue frames represent stem/progenitor cell and luminal cell populations, respectively. **b**-**d** Frequency (**b**, **d**) and size (**c**) of spheres formed by CD49f^hi^/Sca-1^+^ stem/progenitor cells (**b**, **c**) and CD49f^lo^/Sca-1^-^ (**d**) luminal cells from 3-month-old WT, *Mme^-/-^*, *Pten*^PE-/-^, and *Mme^-/-^Pten*^PE-/-^ mice (n=6 per genotype). **e**, **f** Relative frequency of sphere formation by CD49f^hi^/Sca-1^+^ stem/progenitor cells (**e**) and CD49f^lo^/Sca-1^-^ luminal cells (**f**) isolated from 3-month-old WT, *Mme^-/-^*, *Pten^loxP/loxP^*, and *Mme^-/-^ Pten^loxP/loxP^* mice followed by Ad-*Cre* infection (n=6 per genotype). Spheres counts were normalized to the Ad-*blank*-infected spheres of each passage. P0-P3, passages 0-3. **P<0.01, ***P<0.001. Error bars denote SD. **a**-**f** Data represent three independent experiments.

Formation of prostaspheres is used as a functional cell culture test for presence, growth and self-renewal potential of prostate stem/progenitor cells. Consistent with FACS results, CD49f^hi^/Sca-1^+^ cells isolated from prostates of *Mme^-/-^Pten*^PE-/-^mice showed the highest frequency of prostaspheres in multiple consecutive sphere dissociation and regeneration passages (Fig. 4b). Furthermore, prostaspheres deficient for both genes had larger size, as compared to WT, MME or PTEN deficient stem/progenitor cells (Fig. 4c). In all groups CD49f^lo^/Sca-1^-^ luminal cells formed very few spheres after the first plating and no spheres were observed after the first passage (Fig. 4d). Thus, it is unlikely that the increase in prostaspheres in PTEN and MME deficient cells resulted from a reprograming of differentiated cells towards a stem cell state.

To directly test if the observed results represent direct effects of MME and/or PTEN on prostate stem/progenitor cells we isolated CD49f^hi^/Sca-1^+^ stem/progenitor cells and CD49f^lo^/Sca-1^-^ luminal cells from prostates of WT, *Mme*^-/-^, *Pten^loxP/loxP^*, and *Mme*^-/-^*Pten^loxP/loxP^* mice and infected them with adenovirus expressing Cre recombinase (Ad-*Cre*). Consistent with our previous experiments, lack of both *Pten* and *Mme* had the most pronounced effect on frequency of CD49f^hi^/Sca-1^+^ stem/progenitor cells in consecutive passages (Fig. 4e). Luminal cells formed only few spheres with the same frequency in all groups (Fig. 4f). Taken together, these results showed that MME cooperates with PTEN in regulation of prostate stem/progenitor cell functions.

### GRP promotes activities of PTEN-deficient mouse prostate stem/progenitor cells

To identify mechanisms by which MME may affect regulation of prostate stem/progenitor cells, we next examined the expression of MMEs main substrates, GRP, neurotensin (NT) and vasoactive intestinal peptide (VIP) in the prostates of WT, *Mme^-/-^*, and *Pten*^PE-/-^ and *Mme^-/-^Pten*^PE-/-^ strains. Strong GRP expression was detected only in prostates of *Mme^-/-^Pten*^PE-/-^ mice, while NT and VIP were not detected in all cases (Supplementary Fig. 5).

To test if GRP could recapitulate the effects of MME deficiency, we isolated prostate cells from *Mme^-/-^Pten*^PE-/-^ mice, and either performed knockdown of GRP receptor (GRPR, Fig. 5a) or administered GRPR antagonist ([Tyr^4^, D-Phe^12^]-Bombesin. Both approaches reversed effects of MME deficiency on size and formation frequency of prostaspheres (Fig. 5b and 5c).

**Fig. 5.**
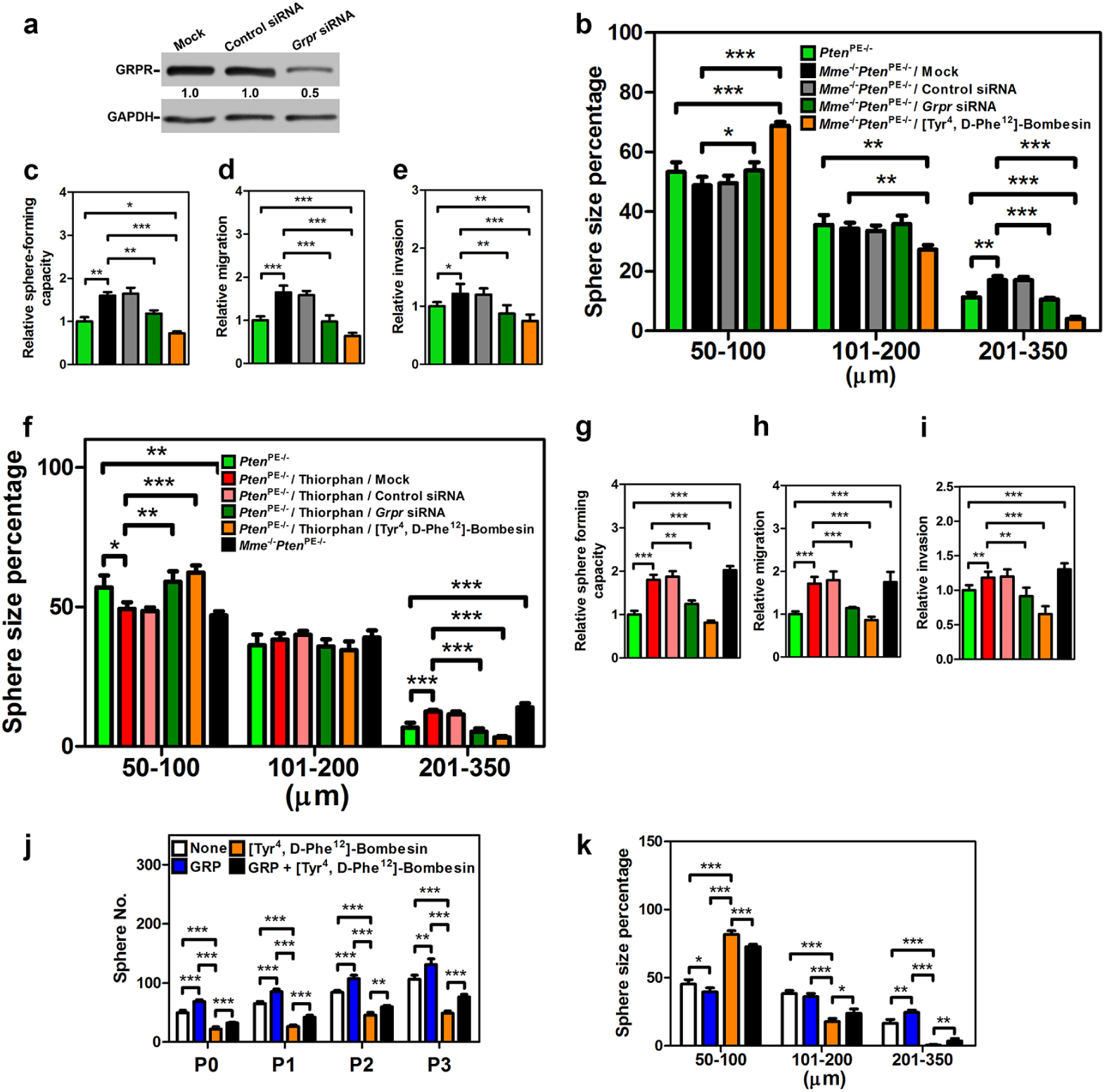
GRP promotes *Pten*-deficient mouse prostate stem/progenitor cell expansion and sphere-forming capacity. **a-e** Western blot of GRPR expression (**a**), prostasphere size (**b**), sphere-forming capacity (**c**), migration (**d**), and invasion (**e**) of prostate cells isolated from 3-month-old *Pten*^PE-/-^ and *Mme^-/-^Pten*^PE-/-^ mice (n=6 per genotype) were performed. **f**-**i** GRP recapitulates the effects of MME catalytic inhibition. Prostasphere size (**f**), sphere-forming capacity (**g**), migration (**h**), and invasion (**l**) of prostate cells isolated from 3-month-old *Pten*^PE-/-^ and *Mme^-/-^Pten*^PE-/-^ mice (n=6 per genotype) were performed. *P<0.05, **P<0.01, ***P<0.001. All error bars denote SD. (**j**, **k**) Frequency (**j**) and size (**k**) of prostaspheres formed by Ad-*Cre*-infected CD49f^hi^/Sca-1^+^ stem/progenitor cells isolated from 3-month-old *Pten^loxP/loxP^* mice (n=6) with treatments of GRP and/or [Tyr^4^, D-Phe^12^]-Bombesin were performed. P0-P3, passages 0-3. Annotations for groups in **f**-**l** are shown in **f**. Annotations for groups in **j** and **k** are shown in **j**. HE, hematoxylin and eosin staining *P<0.05, **P<0.01, ***P<0.001. All error bars denote SD. **a**-**g** Data represent three independent experiments.

Consistent with the observation of frequent vascular invasion by prostate adenocarcinomas in *Mme^-/-^Pten*^PE-/-^ mice, we detected increased cell motility and invasion of prostate cells isolated from *Mme^-/-^Pten*^PE-/-^ mice, as compared to those prepared from *Pten*^PE-/-^ mice (Fig. 5d and 5e). This effect was reversed by either GRPR knockdown or treatment with [Tyr^4^, D-Phe^12^]-Bombesin. We next isolated prostate cells form *Pten*^PE-/-^ mice, and treated them with the MME enzyme inhibitor Thiorphan. Sphere size (Fig. 5f), sphere forming capacity (Fig. 5g), cell motility (Fig. 5h), and invasion (Fig. 5l) were each increased by MME catalytic inhibition; and more importantly *Grpr* siRNA knockdown or a GRPR antagonist abrogated the stimulated functions of MME enzyme inhibition on the prostate cells form *Pten*^PE-/-^ mice.

Finally, to test if GRP directly affects the function of PTEN-deficient prostate stem/progenitor cells, CD49f^hi^/Sca-1^+^ prostate stem cells isolated from *Pten^loxP/loxP^* were infected with Ad-*Cre* followed by treatment with GRP and/or [Tyr^4^, D-Phe^12^]-Bombesin. GRP addition reproduced the effects of MME deficiency on formation frequency and size of prostasphere, while these effects were abrogated by [Tyr^4^, D-Phe^12^]-Bombesin (Fig. 5j and 5k). Taken together, these data suggest that the MME substrate GRP mediates in part the growth and invasive effect of PTEN-deficient prostate cells observed in association with loss or catalytic inhibition of MME; and GRP is a key MME target responsible for stimulating prostate stem/progenitor cells.

### GRP promotes expansion of human prostate cancer propagating cells

To further assess the relevance of our observations to human disease we tested the effects of GRP and [Tyr^4^, D-Phe^12^]-Bombesin on PTEN-deficient androgen-refractory human prostate cancer cells DU145 and PC3. Consistent with previous reports ^34, 41, 48^, GRP promoted cell proliferation, migration, and invasion, and these effects were negated by addition of [Tyr^4^, D-Phe^12^]-Bombesin in both cell lines (Supplementary Fig. 6). Similar effects cell migration and invasion were also observed in androgen-sensitive human prostate adenocarcinoma cells LNCaP (Supplementary Fig. 7). In agreement with our observations on mouse prostate epithelium cells, GRPR knockdown rescued the stimulating effects of GRP and inhibitory effects of [Tyr4, D-Phe12]-Bombesin, suggesting the crucial role of GRP/GRPR in control of the above parameters in both species (Supplementary Fig. 6 and 7). GRP treatment also increased expression of synaptophysin in DU145, PC3 cells and LNCaP cells, thereby suggesting that GRP/GRPR signaling may influence neuroendocrine commitment (Supplementary Fig. 8).

Next we tested if GRP/GRPR signaling has an immediate effect on prostate cancer cells with stem cell-like properties. Such cells, called cancer propagating or cancer stem cells ^13, 18^, are characterized by their ability for long-term self-renewal, high potential for proliferation, high tumorigenicity, capacity to generate the whole spectrum of heterogeneity of their original tumors, and resistance to androgen withdrawal. Prostate cancer propagating cells can be isolated by either high enzymatic activity of ALDH (ALDEFLUOR assay) ^22^ or by detection of CD44 expression ^33^. In both approaches cancer propagating cells isolated from DU145, PC3 and LNCaP cells were increased in number after GRP treatment (Fig. 6a-6d and Supplementary Fig. 7e and f). Cancer propagating cells also formed spheres in greater number and size. These changes were diminished by [Tyr^4^, D-Phe^12^]-Bombesin and/or GRPR siRNA (Fig. 6a-6g).

**Fig. 6.**
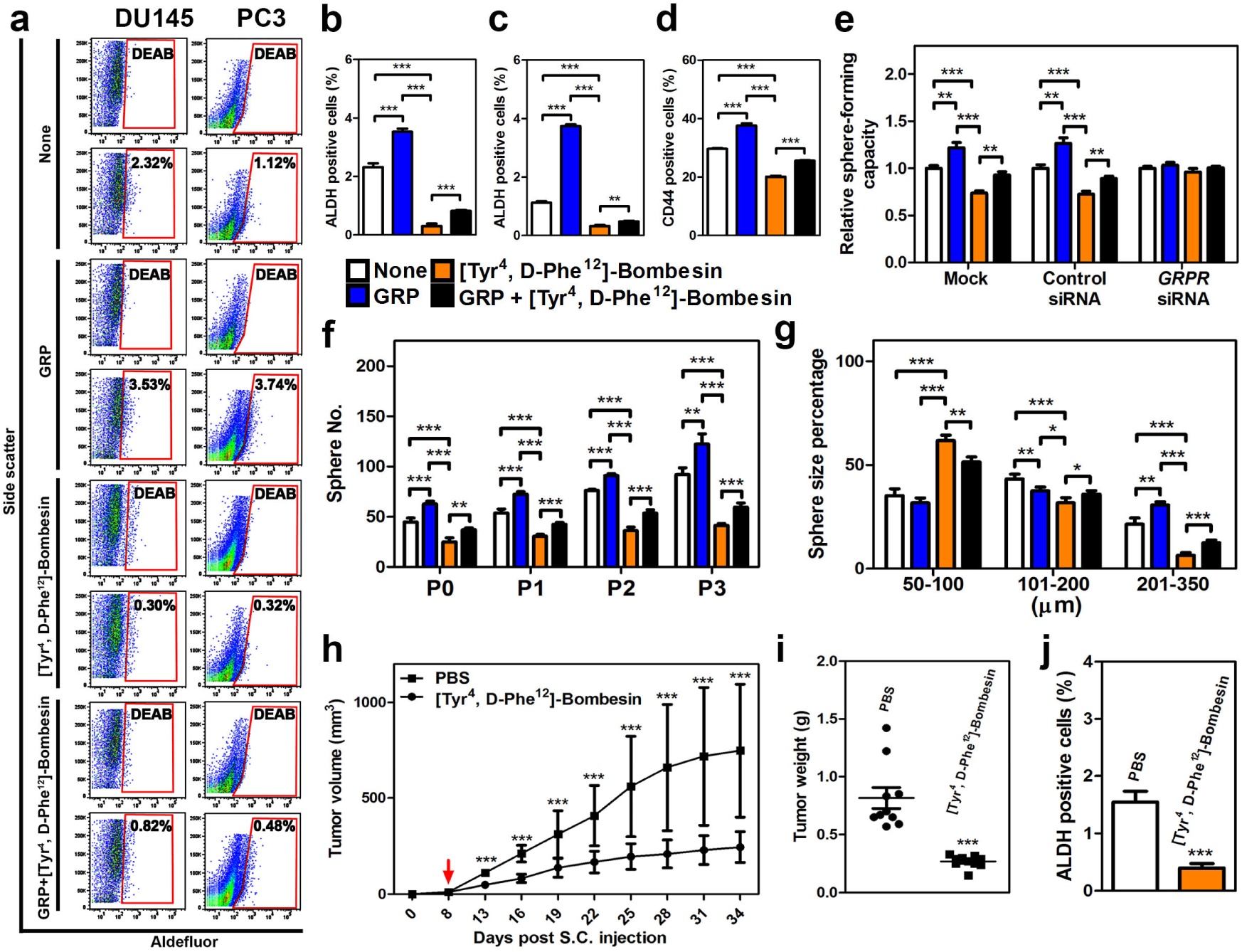
GRP promotes activities of human prostate cancer propagating cells. **a-e** ALDEFLUOR assay (**a**), quantitative analysis of ALDH positive cancer propagating cells (%) of DU145 (**b**) and PC3 (**c**), CD44 positive cancer propagating cells (%) of DU145 (**d**), and sphere-forming capacity of DU145 (**e**) with treatment of GRP and/or [Tyr^4^, D-Phe^12^]-Bombesin were performed. (**a**) The number in each red frame represents the percentage of ALDH positive cancer propagating cells. **f** and **g** Frequency (**f**) and size (**g**) of prostaspheres formed by FACS-purified CD44 positive cancer propagating cells of DU45 with treatments of GRP and/or [Tyr^4^, D-Phe^12^]-Bombesin are shown. P0-P3, passages 0-3. **h** Average volume of PC3 tumor xenografts at indicated time points. Red arrow indicates the starting day of i.p. injection of [Tyr^4^, D-Phe^12^]-Bombesin or PBS. **i** Average weight of PC3 tumor xenografts at sacrifice (day 34). **j** ALDH positive cancer propagating cells (%) of PC3 tumor xenografts. *P<0.05, **P<0.01, ***P<0.001. All error bars denote SD. Data represent three independent experiments (**a**-**g**), and two groups of five mice with injection of [Tyr^4^, D-Phe^12^]-Bombesin or PBS, respectively (**h**-**j**).

To study the effect of GRP on human prostate cancer xenografts *in vivo*, immunodeficient NSG mice were subcutaneously injected with PC-3 cells and treated with [Tyr^4^, D-Phe^12^]-Bombesin when the tumor size reached 10 mm^3^ (Fig. 6h-6j). Daily intraperitoneal injections of [Tyr^4^, D-Phe^12^]-Bombesin reduced both tumor volume and weight by 70% by the time of animal euthanasia at 26 days after the beginning of treatment. Importantly, according to the ALDEFLUOR assay, a fraction of prostate cancer propagating cells was significantly reduced in tumors treated with [Tyr^4^, D-Phe^12^]-Bombesin as compared to controls treated with vehicle alone. The same outcomes have been observed after transplantation of PC3 cells infected with lentivirus expressing GRPR shRNA (Supplementary Fig. 9). Thus, the effects of GRP/GRPR signaling abrogation in inhibiting prostate cancer cell tumorigenicity result from diminishing the population of cancer propagating cells.

## Discussion

Our study provides direct genetic support to previous reports proposing *Mme* role as a tumor suppressor gene. Our findings show that *Mme* cooperates with *Pten* in suppression of prostate carcinogenesis. Lack of *Mme* leads to formation of advanced adenocarcinomas characterized by intravascular invasion, a feature not observed adenocarcinomas of *Pten*^PE-/-^ mice. In agreement with their more aggressive behavior adenocarcinomas of composite mice also have higher proliferative rate, increased number of CK5 and p63 positive cells and elevated levels of EZH2 expression.

It has been reported previously, that *Pten* conditional deletion in the prostate epithelium leads to basal cell proliferation with concomitant expansion of the prostate stem/progenitor-cell like Sca-1^+^ and BCL-2^+^ subpopulation ^51^. CD49f^hi^/Sca-1^+^ *Pten*- deficient neoplastic cells exhibits cancer propagating cell properties, such as high sphere-forming capacity, sustained self-renewal, increased proliferation, and tumorigenic potential, as compared with other isogenic subpopulations ^9, 28^. According to our studies, the combined *Mme* and *Pten* deficiency leads to further increase of number and growth potential of prostate stem/progenitor cells. These observations suggest that *Mme* and *Pten* cooperate in controlling stem/progenitor cells. However, since lack of *Pten* may also stimulate a basal to luminal transdifferentiation in conjunction with proliferation ^6, 20, 52^, further studies are needed to evaluate the role of cell plasticity as an additional factor influencing effects of combined *Mme* and *Pten* deficiency. Our findings also show that the lack of both *Mme* and *Pten* leads to the formation of dysplastic lesions and adenocarcinomas in the proximal regions of prostatic ducts, the structures particularly enriched in stem/progenitor cells. Thus *Mme* may playing a particularly critical role at the site of prostate stem/progenitor cell niche.

Our findings also show that effects of MME on prostate stem/progenitor cells depend on the presence of its downstream effector, GRP, Furthermore, we have observed that abrogation of GRP/GRPR signaling may diminish the pool of cancer propagating cells, thereby highlighting an important role of neuroendocrine signaling in the regulation of cell stemness. Previously, prostate cancer propagating cells have been shown to be regulated by the PTEN/PI3K/AKT pathway ^9^. Our study suggests that this effect is induced by neuropeptide signaling regulated by MME and is potentiated by MME loss. This provides a rationale for using the MME/GRP pathway for targeting of cancer propagating cells, perhaps as a part of combinatorial therapy approaches.

Our study shows that accelerated progression of prostate carcinomas associated with MME deficiency is not associated with increase in the number of neuroendocrine cells. Neuroendocrine cells are important for prostate development ^4^. The majority of human prostate adenocarcinomas and mouse models associated with *Pten* deficiency contain neuroendocrine cells ^23, 58^. However, with the exception of highly aggressive neuroendocrine/small cell carcinomas, it remains debatable if increased number of neuroendocrine cells, as defined by their expression of neuroendocrine markers, such as synaptophysin and calcitonin gene-related peptide, are essential for the accelerated progression of pre-castrate prostate adenocarcinoma ^1, 2, 45^. Our study suggests that accumulation of un-cleaved neuropeptides may mitigate the requirement for neuroendocrine cell expansion or lead to a feedback inhibition of neuroendocrine cell differentiation in MME-deficient prostate cancers.

The recent introduction of therapies that better target the androgen axis has led to a significant increase in frequency of castrate-resistant prostate cancer with neuroendocrine differentiation ^8^. It has been shown that inactivation of the *RB1* tumor suppressor gene increases cell plasticity along with the promotion of an aggressive neuroendocrine phenotype ^19, 27, 32^. In castrated mice with combined *Pten* and *Trp53* deficiency some prostate tumors arising from luminal cells had regions of highly proliferative cells with overt neuroendocrine differentiation. However, other tumors showed only limited foci of neuroendocrine differentiation without any detectable proliferation ^58^. In this report *Pten* inactivation alone resulted in modest increase in neuroendocrine foci, consistent with our findings. In our model associated with *Pten* and *Mme* deficiency we observed increased number of neuroendocrine neoplastic clusters and increased proliferation in such clusters after castration. Furthermore, as compared to *Pten*^PE-/-^ mice, recurrent tumors in castrated *Mme^-/-^Pten*^PE-/-^ mice had more aggressive phenotype based on increased levels of phosphorylated MET, and reduced expression of E-cadherin. The mechanisms by which MME deficiency facilitates neuroendocrine trans-differentiation of the prostate epithelium or leads to expansion neuroendocrine cell lineage after castration remain to be investigated. It also remains to be investigated if MME deficiency represents an alternative to *RB1* loss for progression of some castrate-resistant prostate cancer. Our mouse model based on inactivation of *Mme* and *Pten* should offer an important tool for answering these important questions of prostate cancer pathogenesis and testing new therapeutic approaches.

## Materials and Methods

### Mice

*ARR2PB-Cre* transgenic male mice on FVB/N background (*PB-Cre4*) ^53^ were crossed with *Pten^loxP/loxP^* ^21^ female mice on the 129/BALB/c background. Resulting *PB-Cre4Pten^loxP/loxP^* male mice were designated as *Pten*^PE-/-^ mice. *Pten*^PE-/-^ male mice were crossed with *Mme* null female mice on C57BL6 background (*Mme^-/-^*) ^24^. Offspring with *PB-Cre4 Mme^-/-^ Pten^loxP/loxP^* genotype were designated as *Mme^-/-^Pten^PE-/-^* mice. To minimize the confounding effects of genetic background *Pten*^PE-/-^ and *Mme^-/-^Pten*^PE-/-^ mice were backcrossed to FVB/N for at least 10 crosses and all control experiments were performed on age and sex-matched randomized mice of the same background. All animal experiments were carried out in strict accordance with the recommendations of the Guide for the Care and Use of Laboratory Animals of the National Institutes of Health. The protocol was approved by the Institutional Laboratory Animal Use and Care Committee at Cornell University. All efforts were made to minimize animal suffering.

### Immunohistochemistry and quantitative image analysis

Immunoperoxidase staining of paraffin sections of paraformaldehyde-fixed tissue was performed by a modified Elite avidin-biotin-peroxidase (ABC) technique ^5^. Antigen retrieval was done by boiling the slides in 10 mM citric buffer (pH 6.0) for 10 min. The primary antibodies to MME (Santa Cruz; Dallas, TX; #sc-80021, 1:100), Ki67 (Leica Microsystems; Bannockburn, IL; #NCLKi67p, 1:1000), EZH2 (Cell Signaling Technology; Danvers, MA; #5246, 1:200), keratin 5 (CK5, Covance; Dallas, TX, #PRB-160P, 1:2500), keratin 8 (CK8, Developmental Studies Hybridoma Bank; Iowa City, IA; #TROMA-I, 1:10), p63 (Santa Cruz, #sc-8431, 1:1000), PTEN (Cell Signaling Technology, #9559S, 1:800), pAKT (Cell Signaling Technology, # 3787S, 1:50), SYP (BD Biosciences; #611880, 1:500), GRP (Santa Cruz, #sc-7788, 1:100), NT (Santa Cruz, #sc-20806, 1:1000), and VIP (Abcam; Cambridge, MA; #ab8556, 1:200), E-cadherin (Cell Signaling Technology #3195S, 1:200), phospho-MET (Cell Signaling Technology, #3077, 1:50), were incubated with deparaffinized sections at 4°C overnight, followed by incubation with secondary biotinylated antibody (1 hour, room temperature) and modified avidin-biotin-peroxidase (ABC) technique. Methyl green was used as the counterstain in immunoperoxidase stainings. Slides were scanned by ScanScope CS (Leica Biosystems, Vista, CA) with 40X objective followed by lossless compression. For double fluorescence antibodies against SYP (BD biosciences, #611880) and Ki67 (abCAM, #16667) were used at a concentration of 1:40 and 1:50 respectively, for overnight incubation at 4°C, followed by the incubation with secondary antibodies (Alexa Fluor 568 donkey-anti-mouse #A10037, Alexa Fluor 488 donkey-anti-rabbit #21206, Life Technologies, 1:200 each, 2 hours, room temperature) and counterstaining with 4’6-diamidino-2-phenylindole (DAPI, Sigma-Aldrich #D9542). Immunofluorescent sections were observed and imaged using Leica TCS SP2 confocal laser scanning microscope system (Leica Microsystems). Quantitative analysis of immunohistochemistry (IHC) was performed with the ImageJ software (W. Rasband, National Institutes of Health, Bethesda, MD).

### Tumorigenicity experiments

Human prostate cancer PC3 cells (5 x 10^6^ cells) were suspended in the mixture of 200 µl PBS and 200 µl Matrigel (BD Biosciences, #356237) and injected subcutaneously into 5 weeks old NSG (NOD.Cg-*Prkdc*^scid^*Il2rg*^tm1Wjl^/SzJ4; The Jackson Laboratory, stock number #005557) male mice. Intra-peritoneal injection of GRPR antagonist (0.8 μg/g body weight/day in PBS) was started when the tumor size reached 10 mm^3^. After 26 days of treatment with GRPR antagonist, mice were euthanized and subjected to necropsy. Tumor xenografts were collected pathological evaluation and ALDEFLUOR assay followed by FACS analysis.

### Castration experiments

Male mice were weighed and anesthetized using isoflurane. Pressure was applied to the abdomen of each mouse to push both testes down into the scrotal sac. An approximately 1 cm incision was made across the midline of the scrotal sac to expose the testicular membrane. The midline between the left and right testes sacs were located and a small incision is made on the left side of the membrane midline. The testes, attached fatty tissue, the vas deferens and associated blood vessels are carefully pulled out through the incision. The blood vessels were cauterized and the testes was removed by severing below the cauterization. Same procedure was repeated to remove testes from the right side of the midline. Remaining tissue was pushed back into incision which was then closed with wound clips. Post procedure, the mouse was placed on a heating pad till it recovered from the effects of the anesthetic followed by placement in a clean cage with fresh chow and water.

### Pathologic assessment

Moribund mice, as well as those sacrificed according to schedule, were anesthetized with avertin and, if necessary, subjected to cardiac perfusion at 90 mm Hg with PBS. After macroscopic evaluation during necropsy, lung, liver, prostate, and lymph nodes were fixed in phosphate-buffered 4% paraformaldehyde tissues, embedded in paraffin and 4-μm-thick sections were stained with Hematoxylin (Mayer’s haemalum) and eosin. Striated muscle layer was used for identification of proximal (periurethral) and distal regions of prostatic ducts in transverse sections, as previously described ^57^. Mouse prostatic intraepithelial neoplasia (PIN) and adenocarcinoma were defined according to earlier publications ^5, 16, 31, 56^. Briefly, distal PIN 1 has 1 or 2 layers of atypical cells; PIN 2 has 3 or more layers of atypical cells, PIN 3 occupies the near entire glandular lumen; and PIN 4 fills and distorts the glandular profile, and is frequently marked by pronounced desmoplastic reaction. In agreement with the recent consensus report ^16^, PIN1 and PIN2 represent low-grade PIN (LG-PIN) and PIN3 and 4 represent high-grade PIN (HG-PIN). Due to different architecture of the proximal regions of prostatic ducts, current PIN classification cannot be carefully applied to atypical proliferative lesions found in those structures. Thus we named those lesions as proximal duct dysplasia to stress their dissimilarity to PIN of distal regions of prostatic ducts. Given the complexity and controversial nature of the interpretation of stromal microinvasion we used term early adenocarcinoma only for neoplasms with invasive stromal growth confirmed by serial sections followed by 3D reconstruction. We used term advanced adenocarcinoma for neoplasms invading blood and lymphatic vessels. All pathological evaluations were performed in blinded fashion.

### Cell culture

DU145, PC3 and LNCaP cell lines were obtained from the American Type Culture Collection (ATCC) and cultured in minimum essential medium (Cellgrow, #10-010-CV), F-12K (Cellgrow, #10-025-CV) and RPMI 1640 (VWR, cat # 4500-396), respectively, supplemented with 10% heat-inactivated fetal bovine serum (FBS, GIBCO, #16141-079) and Penicillin-Streptomycin (Cellgrow, #30-002-Cl). The cultures were maintained at 37°C in a 5 % CO2 incubator. All cell lines were confirmed to be free of mycoplasma.

Primary mouse prostate cells were isolated following described procedures ^5, 25^. For gastrin-releasing peptide (GRP) experiments, 5 nM GRP (Sigma, #G8022) and 4 μM GRPR antagonist ([Tyr^4^, D-Phe^12^]-Bombesin, Sigma, #B0650) were used for cell culture experiments. Media was changed every 3 days. Lipofectamin 2000 reagent (Invitrogen; Carlsbad, CA, #11668-030) was used for the transfection following manufacturer’s recommendations. Control siRNA (Santa Cruz, #sc-37007) and two independent human *GRPR* siRNAs (Santa Cruz, # sc-106924 and Life Technologies, #145216), human GRPR shRNA in Lentiviral Vector (Genomics Online; Limerick, PA, # ABIN3479889) and mouse *Grpr* siRNAs (Life Technologies, #157912 and #157914) were used for all knockdown experiments. MME inhibitor DL-Thiorphan, (1 μM, Sigma, #T6031) ^35^ was used to treat cells at 37°C for 20 min, followed by subsequent experiments.

5-bromo-2’deoxyuridine (BrdU) staining migration and invasion assays were performed as previously described ^5, 11^.

### ALDEFLUOR assay and Fluorescence Activated Cell Sorting

For detection of aldehyde dehydrogenase (ALDH) enzymatic activity, 10^6^ cells were placed in ALDEFLUOR buffer and processed for staining with the ALDEFLUOR Kit (STEM CELL, #01700) according to the manufacture’s protocol. Unstained and ALDH inhibitor (diethylaminobenzaldehyde, DEAB)-treated cells served as controls. For detection of CD44 expressing cancer cells, prostate cancer cells were stained for CD44 (BD Bioscience, #553134) to sort out CD44 positive and CD44 negative cells. Cell sorting and data analysis were performed on a FACS Aria II sorter equipped with the FACS DiVa software (BD Bioscience).

### Prostasphere assay

The preparation of prostate epithelial cell suspensions, stem/basal cells, and luminal cells from male mice were performed based on previously described prostrate sphere assays ^5, 25^. Briefly, 10^4^ mouse prostate stem/basal cells, mouse prostate luminal cells, human prostate cancer cells, and human prostate cancer propagating cells were resuspended in 120 µl of a 1:1 mixture of Matrigel (BD Biosciences, #354234) and PrEGM (Lonza, #CC-3166), and plated around the rim of a well of a 12-well tissue culture plate. Matrigel mix was allowed to solidify at 37°C for 15 min, and 1 ml of PrEGM was added per well. Media was changed every 3 days. To recover the spheres, each well was treated with enzyme mixture: 750 µl Collegenase/Dispase 4 mg/ml (Roche, #10269638001), 30 mg BSA (Sigma, #A3311), and 1 µl DNase1 10mg/ml (Sigma, D4513), followed by Trypsin 0.25% EDTA (Cellgrow, 25-052-Cl) to make cell suspensions, which were ready for passage.

### Quantitative real-time PCR

RNA was extracted using TRIzol reagent according to manufacture’s instructions (Thermo Fisher). cDNA was produced using the SuperScript III First-Strand Synthesis kit (Thermo Fisher). Real-time PCR was performed using PerfeCTa SYBR Green Super Mix Reagent (Quanta bio) on C1000 Touch Thermal Cycler PCR machine (Bio-Rad). *SYP* expression was assessed using forward primer 5’-TGCGCTAGAGCATTCTGGG-3’ and reverse primer 5’-CTTAAAGCCCTGGCCCCTTCT-3’.

### Western blot

For western blot cell lysates were prepared using RIPA buffer (50 mM Tris-HCl, (pH 7.4), 1% Nonidet P-40, 0.25% Na-deoxicholate, 150 mM NaCl, 1 mM EDTA, 1 mM PMSF, Aprotinin, leupeptin, pepstatin: 1 μg/ml each, 1 mM Na3VO4, 1 mM NaF), followed by sonication for 10 seconds 5 times on ice. Lysates were then separated by 12% SDS-PAGE and transferred to PVDF membrane (Millipore #IPVH00010). The membrane was incubated overnight at 4°C with antibodies to detect GRPR (Santa Cruz, #sc-32903, 1:1000), and GAPDH (Advanced Immunohistochemical Inc.; Long Beach, CA; #2-RGM2,1:5000), followed by incubation for 1 hour at room temperature with corresponding horseradish peroxidase-conjugated anti-rabbit secondary antibodies (Santa Cruz, #sc-2004, 1:2000) or anti-mouse secondary antibodies (Santa Cruz, #sc-2005, 1:2000) and developed using chemiluminescent substrate (Thermo Scientific, Rockford, IL, #34077).

### Statistics

Statistical analyses were performed with InStat 3.10 and Prism 7 software. (GraphPad, Inc., San Diego, CA). Two-tailed unpaired t-test, direct Fisher’s tests, and log-rank Mantel-Haenszel test were used as appropriate. To ensure adequate power to detect a pre-specified effect size all sample sizes were chosen based on initial pilot experiments, including animal studies. No samples or animals were excluded from the analysis. The variance was similar between the groups that were statistically compared.

## Acknowledgments

We would like to thank David Dupee, Aditi Iyengar, and Elaina Wang for expert technical assistance, Lavanya Sayam (NYSTEM supported FACS Core) for her help with fluorescence-activated cell sorting and all members of the Nikitin Lab for their advice and support. This work has been supported by NIH (CA096823 and CA197160) and NYSTEM (C023050 and C028125) grants to A.Y.N., NIH (CA72717) and the Genitourinary Oncology Research fund (Weill Cornell) to DMN, and fellowship funding from the Cornell Comparative Cancer Biology Training Program to C.-Y. C.

