## Supplementary Information for "Membrane metalloendopeptidase suppresses prostate carcinogenesis by attenuating effects of gastrin-releasing peptide on stem/progenitor cells"

**Supplementary data**

**Supplementary Table 1. Prostatic Lesions in *Mme*$^{-/-}$, *Pten*$^{PE-/-}$ and *Mme*$^{-/-}$*Pten*$^{PE-/-}$ Mice**

| Age (m) | Prostatic region | Lesion | Strain | | | Fisher's P* |
|---|---|---|---|---|---|---|
| | | | *Mme*$^{-/-}$ | *Pten*$^{PE-/-}$ | *Mme*$^{-/-}$ *Pten*$^{PE-/-}$ | |
| 3 | **Proximal** | None | 100 (3/3)[#] | 100 (16/16) | 83 (5/6) | 0.3043 |
| | | Dysplasia | 0 (0/3) | 0 (0/16) | 17 (1/6) | 0.2727 |
| | | Early adenocarcinoma | 0 (0/3) | 0 (0/16) | 0 (0/6) | NA |
| | | Advanced adenocarcinoma[&] | 0 (0/3) | 0 (0/16) | 0/(0/6) | NA |
| | **Distal** | None | 100 (3/3) | 100 (0/16) | 0 (0/6) | NA |
| | | Low-grade PIN | 0 (0/3) | 6 (1/16) | 0 (0/6) | 1 |
| | | High-grade PIN | 0 (0/3) | 94 (15/16) | 100 (6/6) | 1 |
| | | Early adenocarcinoma | 0 (0/9) | 0 (0/16) | 0 (0/6) | NA |
| | | Advanced adenocarcinoma | 0 (0/9) | 0 (0/16) | 0 (0/6) | NA |
| 7 | **Proximal** | None | 100 (7/7) | 100 (11/11) | 0 (0/8) | NA |
| | | Dysplasia | 0 (0/7) | 0 (0/11) | 87 (7/8) | 0.0002 |
| | | Early adenocarcinoma | 0 (0/7) | 0 (0/11) | 13 (1/8) | 0.4211 |
| | | Advanced adenocarcinoma | 0 (0/7) | 0 (0/11) | 0 (0/8) | NA |
| | **Distal** | None | 100 (7/7) | 0 (0/11) | 0 (0/8) | NA |
| | | Low-grade PIN | 0 (0/7) | 0 (0/11) | 0 (0/8) | NA |
| | | High-grade PIN | 0 (0/7) | 100 (11/11) | 100 (8/8) | NA |
| | | Early adenocarcinoma | 0 (0/7) | 0 (0/11) | 0 (0/8) | NA |
| | | Advanced adenocarcinoma | 0 (0/7) | 0 (0/11) | 0 (0/8) | NA |
| 16 | **Proximal** | None | 100 (15/15) | 100 (0/9) | 0 (0/15) | NA |
| | | Dysplasia | 0 (0/15) | 0 (0/14) | 33 (5/15) | 0.0421 |
| | | Early adenocarcinoma | 0 (0/15) | 0 (0/14) | 47 (7/15) | 0.0063 |

| | | | | | |
|---|---|---|---|---|---|
| | Advanced adenocarcinoma | 0 (0/15) | 0 (0/14) | 20 (3/15) | 0.2241 |
| **Distal** | None | 100 (15/15) | 0 (0/14) | 0 (0/15) | NA |
| | Low-grade PIN | 0 (0/15) | 0 (0/14) | 0 (0/15) | NA |
| | High-grade PIN | 0 (0/15) | 71 (10/14) | 13 (2/15) | 0.0025 |
| | Early adenocarcinoma | 0 (0/15) | 29 (4/14) | 53 (8/15) | 0.2635 |
| | Advanced adenocarcinoma | 0 (0/15) | 0 (0/14) | 33 (5/15) | 0.0421 |

* Fisher's exact test comparing number of lesions in *Pten*$^{PE-/-}$ and *Mme*$^{-/-}$*Pten*$^{PE-/-}$ mice

[#]% (number of mice with lesion out of total number of mice).

[&]Advanced adenocarcinoma is defined as adenocarcinoma with vascular invasion.

**Supplementary Figures**

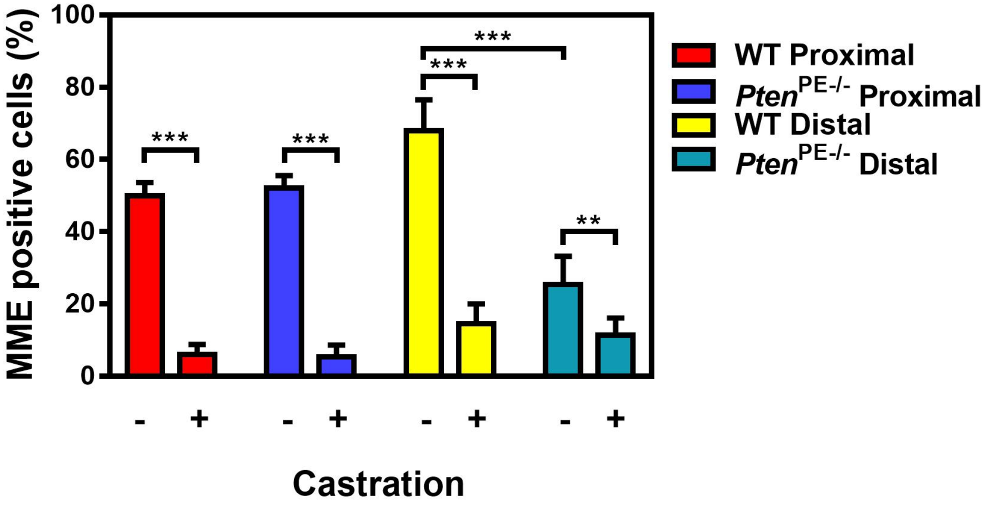


**Supplementary Fig. 1. A quantitative analysis of frequency of MME positive cells in proximal and distal regions of prostatic ducts of wild-type (WT) and *Pten*$^{PE-/-}$ 12 months old mice non-castrated (-) and castrated (+) at 5 months of age.** Areas of stromal invasion by early adenocarcinoma in lack MME expression in *Pten*$^{PE-/-}$ mice and are not included in quantitative analysis. **P<0.01; ***P<0.001. Error bars denote SD. All results are representative of six mice per genotype.

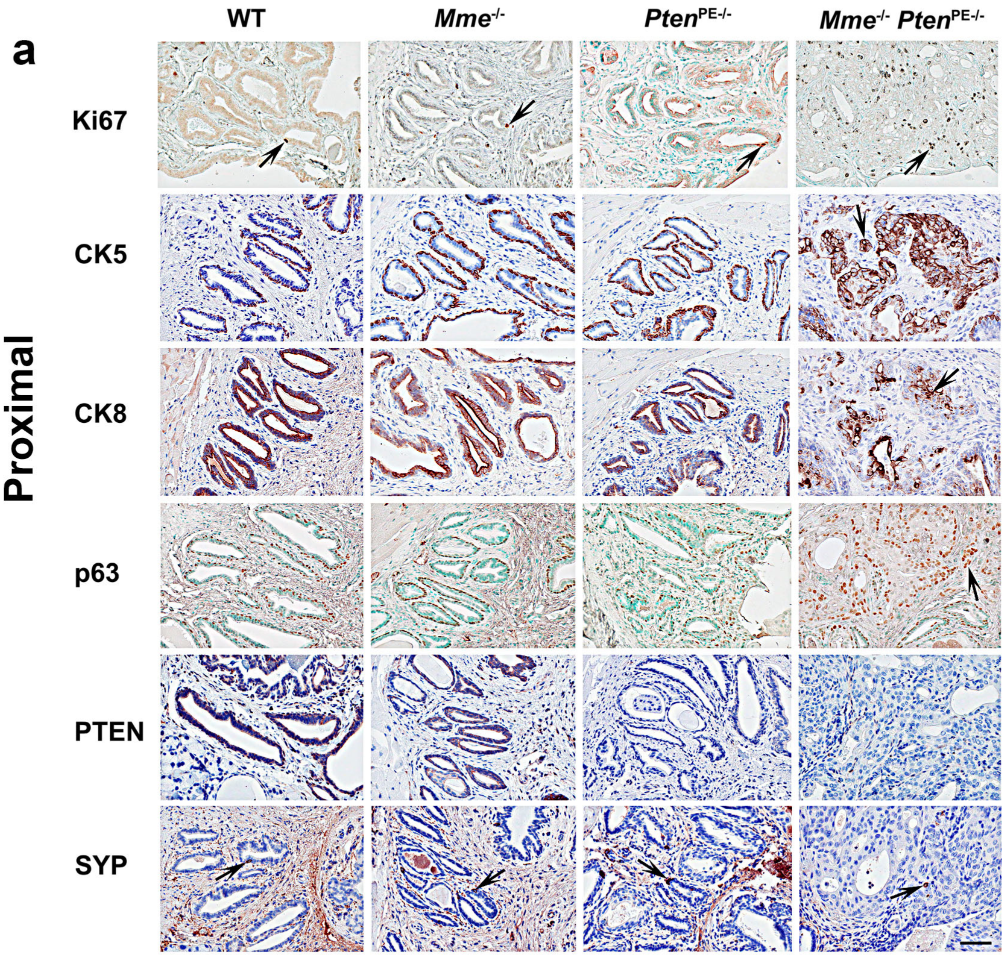
a
WT
Mme-/-
PtenPE-/-
Mme-/- PtenPE-/-
Proximal
Ki67
CK5
CK8
p63
PTEN
SYP

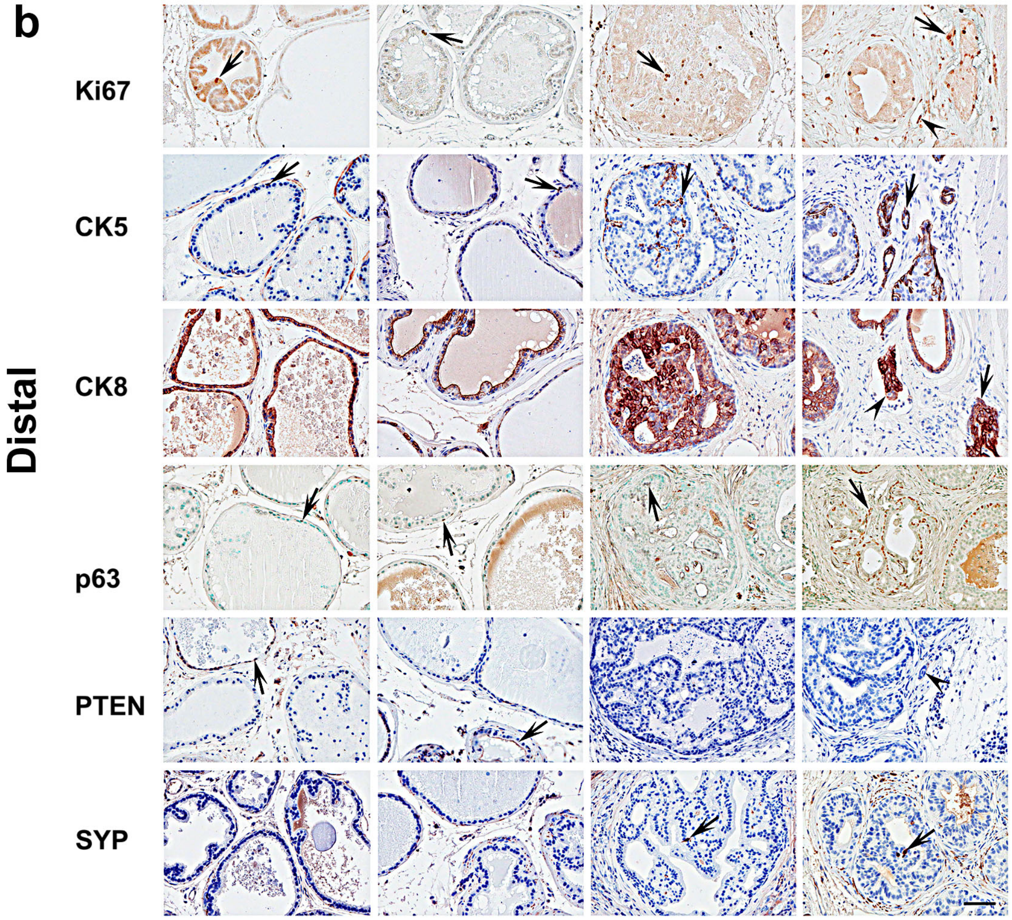


**Supplementary Fig. 2. Characterization of neoplastic lesions associated with PTEN or MME and PTEN deficiency. a-b** Immunostaining of proximal (**a**) and distal (**b**) regions of prostatic ducts. As compared to the prostate epithelium of WT (n=5), *Mme*$^{-/-}$ (n=15) and *Pten*$^{PE-/-}$ (n=14) mice, adenocarcinomas of *Mme*$^{-/-}$*Pten*$^{PE-/-}$ mice (n=15) show an increased number of Ki67, CK5, and p63 positive cells, but no differences in number of synaptophysin (SYP) positive cells. The arrows indicate adenocarcinoma (HE) in *Mme*$^{-/-}$*Pten*$^{PE-/-}$ mice or positive immunostained cells. The arrowheads in the distal region (Ki67, CK8 and PTEN) indicate vascular invasion in *Mme*$^{-/-}$*Pten*$^{PE-/-}$ mice. HE, hematoxylin and eosin staining. The ABC Elite method with hematoxylin (CK5, CK8, PTEN, and SYP) or methyl green (p63) counterstaining was performed. Scale bar, 60 µm for all images.

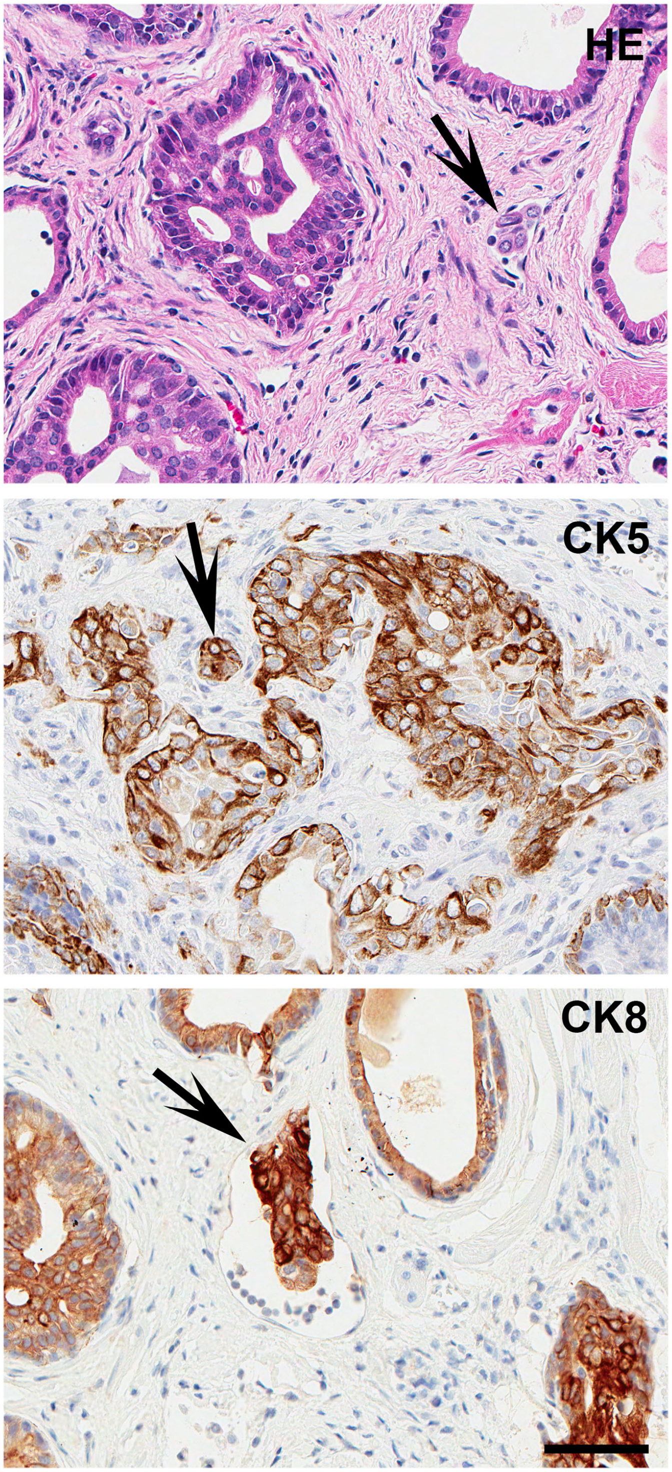


**Supplementary Fig. 3. Vascular invasion of adenocarcinomas in *Mme*$^{-/-}$*Pten*$^{PE-/-}$ mice.** Intravascular carcinoma cells are indicated by arrows. HE, hematoxylin and eosin staining. The ABC Elite method with hematoxylin (CK5, and CK8) counterstaining was performed. Scale bar, 60 μm for all images.

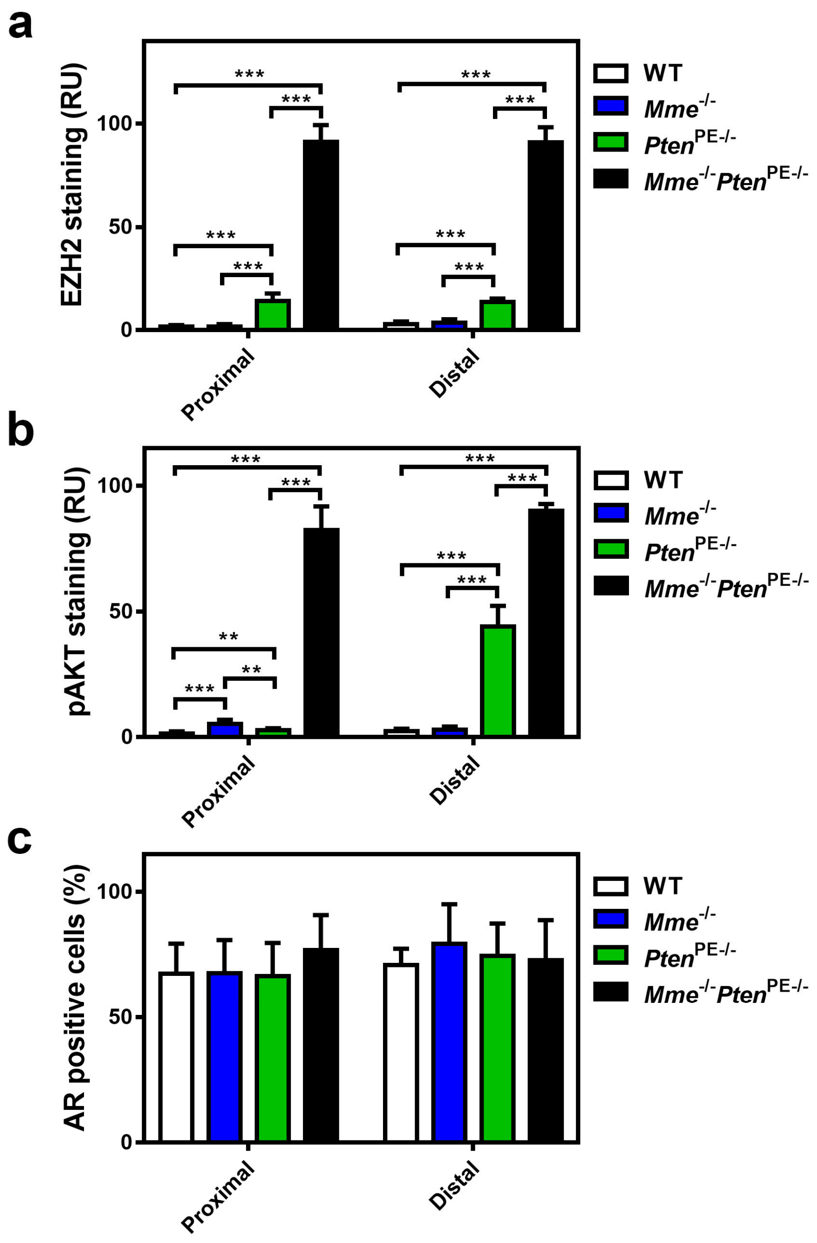


**Supplementary Fig. 4. A quantitative analysis of EZH2 (a), pAKT (b), and AR (c) expression in proximal and distal regions of prostatic ducts.** **P<0.01; ***P<0.001. Error bars denote SD. All results are representative of six mice per genotype.

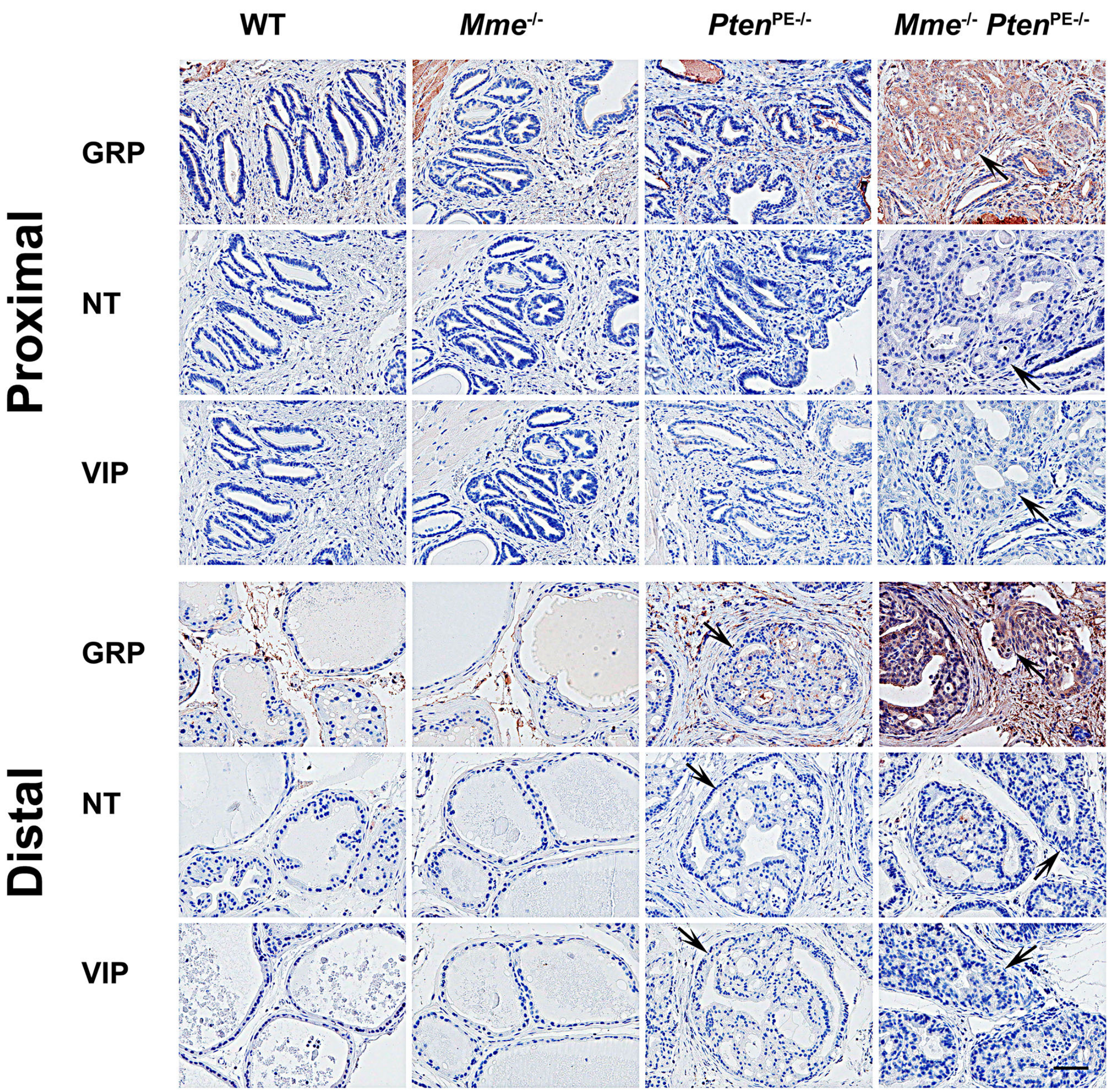


**Supplementary Fig. 5. GRP accumulation in mouse prostatic lesions deficient for *Mme* and *Pten*.** GRP, NT, and VIP expression in the proximal and distal regions of prostatic ducts in 16-month-old WT (n=5), *Mme*$^{-/-}$ (n=15), *Pten*$^{PE-/-}$ (n=14), and *Mme*$^{-/-}$ *Pten*$^{PE-/-}$ (n=15) mice are shown. Arrows, prostatic lesions. The ABC Elite method with hematoxylin counterstaining was performed. Scale bar, 60 μm.

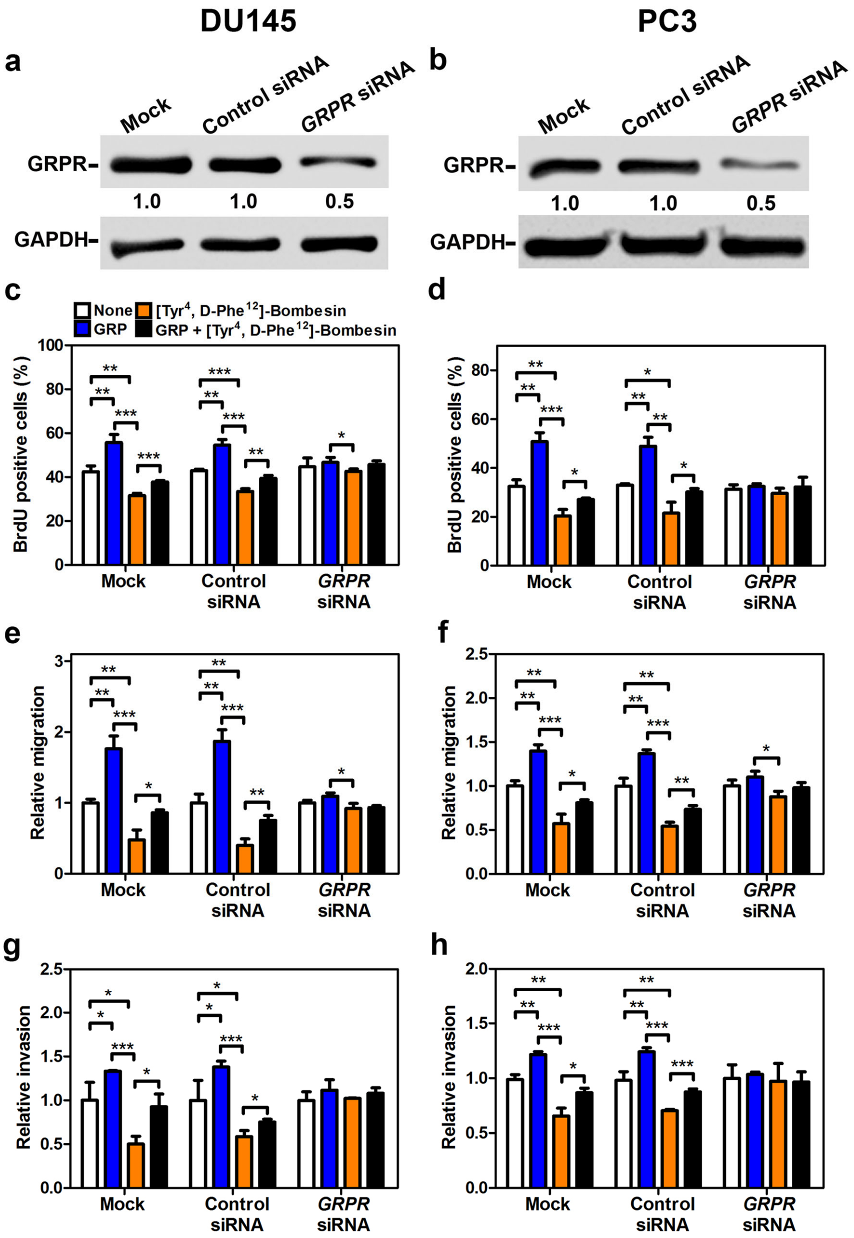


**Supplementary Fig. 6. GRP promotes activities of human prostate cancer cells.** **a**-**h** Western blot of GRPR expression (**a**, **b**) and BrdU positive cells (%) (**c**, **d**), migration (**e**, **f**), and invasion (**g**, **h**) of DU145 (**a**, **c**, **e**, and **g**) and PC3 (**b**, **d**, **f**, and **h**) human prostate cancer cells with treatments of GRP and/or [$Tyr^4$, D-$Phe^{12}$]-Bombesin are shown. *P<0.05, **P<0.01, ***P<0.001. All error bars denote SD. **a**-**h** Data represent three independent experiments.

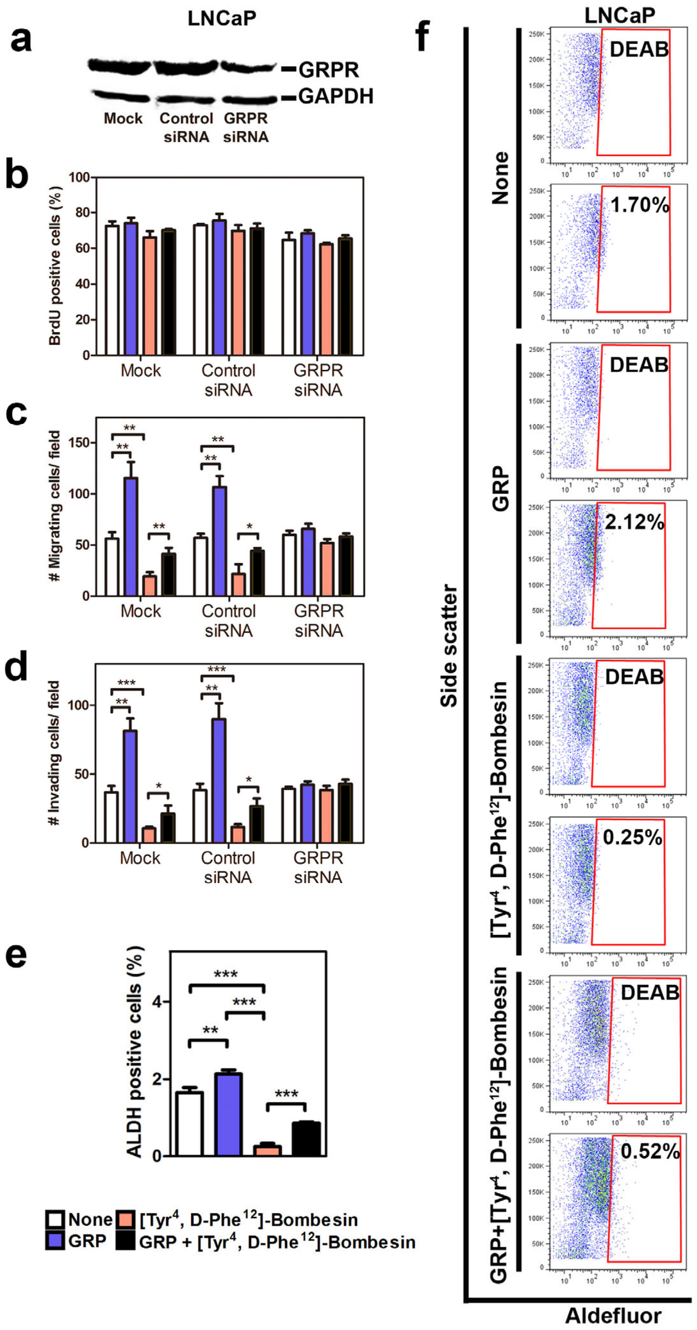


**Supplementary Fig. 7. GRP effects on LNCaP cells. a-d** Western blot of GRPR expression (**a**) and BrdU positive cells (%) (**b**), migration (**c**), and invasion (**d**) of LNCaP cells with treatments of GRP and/or [Tyr$^4$, D-Phe$^{12}$]-Bombesin. **e-f**, quantitative analysis (e) and representative plots of ALDEFLUOR assay for detection of ALDH positive cells (%) in LNCaP cells. *P<0.05, **P<0.01, ***P<0.001. All error bars denote SD. All data represent three independent experiments.

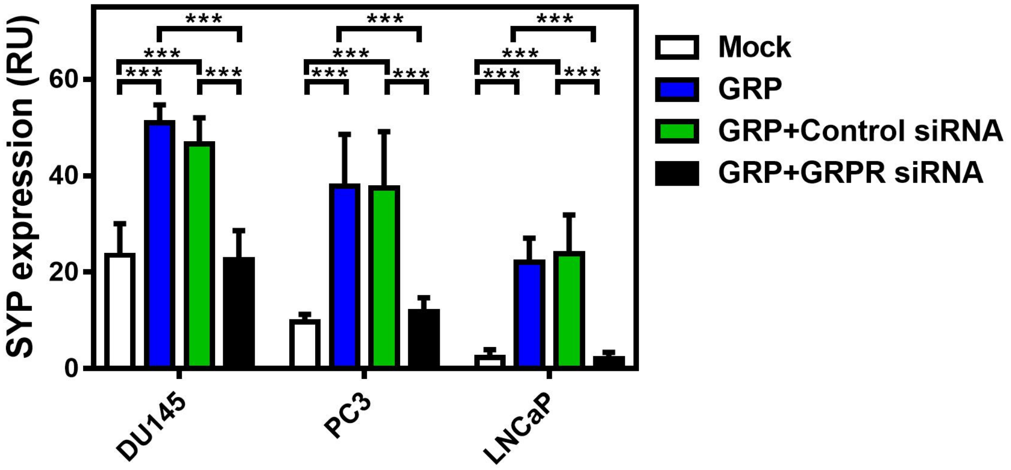


**Supplementary Fig. 8. GRP effects on synaptophysin expression in DU145, PC3 and LNCaP spheres.** ***P<0.001. All error bars denote SD. Data represent three independent experiments.

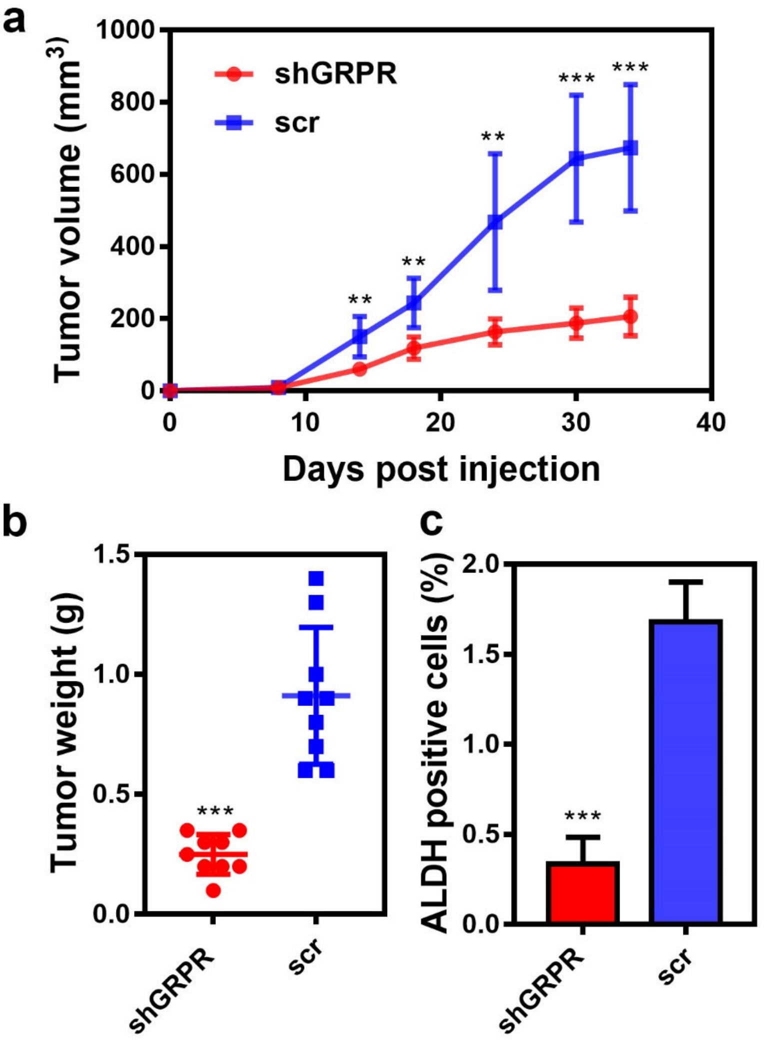


**Supplementary Fig. 9. Effect of GRPR siRNA-mediated knockdown on PC3 tumor xenografts.** **a** Average volume of PC3 tumor xenografts at indicated time points. **b** Average weight of PC3 tumor xenografts at sacrifice (day 34). **c** ALDH positive cancer propagating cells (%) of PC3 tumor xenografts. Data represent two groups of five mice injected with PC3 cells infected with either lenti-shRNA (shCGRP) or lenti-scrambled (scr) control, respectively. *P<0.05, **P<0.01, ***P<0.001. All error bars denote SD.
